# EpiExpr: Predicting gene expression using epigenetic data and chromatin interactions

**DOI:** 10.64898/2026.02.06.704509

**Authors:** Sourya Bhattacharyya, Ferhat Ay

## Abstract

Decoding gene expression from epigenomic landscapes remains a fundamental challenge in genomics. We introduce EpiExpr, a flexible deep learning framework that predicts gene expression from 1D epigenetic tracks (EpiExpr-1D) and integrates 3D chromatin interactions (EpiExpr-3D) to capture distal regulatory effects. Leveraging residual convolutional networks and graph neural networks, including graph attention and graph transformer models, EpiExpr models both local and long-range regulatory influences. Applied to GM12878 and K562 cells, EpiExpr-1D and 3D improve gene expression prediction relative to reference approaches. Analysis using CRISPRi-FlowFISH validated enhancers confirms that EpiExpr-3D accurately prioritizes regulatory elements, compatible with activity-by-contact scores. Remarkably, EpiExpr achieves performance comparable to DNA sequence-based transformer models without requiring sequence embeddings, offering a computationally efficient alternative. This approach provides a scalable, multi-resolution framework (https://github.com/souryacs/3CExpr) for dissecting the contributions of epigenetic modifications and 3D genome organization to gene regulation, enabling broader application across cell types and experimental settings.

## INTRODUCTION

Non-coding DNA, comprising ∼98.5% of the human genome, plays a central role in transcriptional regulation^1^. Epigenomic profiling approaches including ChIP-seq, ATAC-seq, bisulfite sequencing, and DNase-seq^2–4^, together with 3D chromatin contact maps from Hi-C^5,6^, HiChIP^7,8^ and PCHi-C^9^ revealed extensive enhancer–promoter interactions that shape cell-type–specific gene expression^10^. Large-scale consortia such as ENCODE^2,3^ and the 4D Nucleome Project^11,12^ have generated comprehensive datasets characterizing chromatin accessibility, DNA methylation, histone modifications and chromatin architecture, enabling major advances in regulatory genomics. Yet, interpreting these signals and validating regulatory principles across biological contexts remain substantial challenges.

Deep learning has recently emerged as a powerful framework for modeling epigenetic regulation^13^. Convolutional and transformer architectures have been applied to identify regulatory sequence motifs^14^, quantify mutational impacts^15^, simulate epigenomic profiles^16^, model epigenetic signals from DNA sequences^17–19^, predict multi-omic signals^18^, and generate 3D chromatin contacts^20,21^. Transformer-based^22^ foundation models trained on a large number of DNA sequence datasets are used together with explainable-AI techniques^23,24^ to decode the contribution of regulatory sequence motifs^14^, or regulatory regions in transcriptional regulation^25^.

Several models predict gene expression directly from DNA sequence and epigenetic profiles including DeepSEA^26^, Basenji^15^, Enformer^27^, CREaTor^28^, Borzoi^29^ and AlphaGenome^30^ which leverage convolutional neural network (CNN) or transformer to predict gene expression or RNA-seq coverage across cell types. More recent approaches such as EPInformer^31^ incorporate DNA sequence, 1D epigenomic tracks, and 3D chromatin interactions within hybrid CNN + transformer frameworks. Despite their predictive power, sequence-based models are constrained by computational limits that restrict input windows to 200 kb (Enformer) or 524 kb (Borzoi), limiting their ability to capture distal enhancer contributions^32^. Although the recent approach AlphaGenome^30^ considers 1 Mb sequence window, it incurs severe computational complexity and TPUs for processing. Given the high resource demands of transformer architectures, alternative statistical and machine-learning methods have focused on predicting gene expression directly from epigenetic features. Examples include the Activity-By-Contact (ABC) model^33,34^, which integrates enhancer and promoter activity with chromatin contact frequency; 3D-HiChAT, which uses random forests with 1D and 3D epigenomic features^35^; and Epi-GraphReg^36^, which employs epigenetic signals and 3D chromatin interactions to predict gene expression using a combination of CNN and graph attention network (GAT). Although Epi-GraphReg is more scalable due to its architecture, it is limited in using only one cell type at a time, fixed numbers of epigenetic tracks, and fixed track resolutions.

To address these limitations, we introduce **EpiExpr-1D** and **EpiExpr-3D**, two new deep learning frameworks that use residual CNNs, graph attention network^37,38^, and graph transformer architectures (masked label prediction)^39^ to predict gene expression solely from 1D epigenetic tracks (EpiExpr-1D) and also incorporate cell-type–specific 3D Hi-C or HiChIP interactions (EpiExpr-3D). To accommodate the heterogeneity of available epigenomic data across cell and tissue types, we also provide open-source Snakemake pipelines enabling flexible dataset construction with user-defined numbers of cell types, variable resolution epigenetic tracks and CAGE (expression) data. Across benchmarks, both models outperform Epi-GraphReg and achieve predictive accuracy comparable to transformer-based sequence + epigenomic data centric models such as EPInformer, while requiring substantially lower computational resources. Given their high predictive performance, computational efficiency, and broad adaptability, EpiExpr-1D and EpiExpr-3D offer practical, scalable tools for gene-expression modeling across diverse cell types and experimental settings.

## RESULTS

### EpiExpr-1D: A residual CNN model for predicting gene expression from 1D epigenomic profiles

To model gene expression from epigenomic features, we first developed EpiExpr-1D, a deep learning framework that uses only 1D epigenetic profiles (ChIP-seq, ATAC-seq, DNase-seq, etc.) as input. In contrast to Epi-GraphReg^36^, which trains on a single cell or tissue type and uses a fixed set of epigenetic tracks, EpiExpr-1D flexibly supports one or more cell types and accommodates varying numbers and types of available epigenomic datasets for each cell type, similar in spirit to multi-cell-type models such as Enformer^27^.

We implemented an open-source Snakemake pipeline (https://github.com/souryacs/3CExpr/tree/main/EpiExpr_1D_Create_Training_Data) to construct training and validation datasets for *K* ≥ 1 cell or tissue types, each with *N_k_* epigenetic tracks plus a corresponding CAGE track (**Fig. 1A; Methods**). Unlike Epi-GraphReg, which is restricted to fixed resolutions of 100 bp (epigenetic) and 5 kb (CAGE), EpiExpr-1D allows user-defined resolutions for both track types, denoted *e* and *c*, under the constraints that each dataset uses a single pair (*e*, *c*) and that *c* is an integer multiple of *e*. Similar to Epi-GraphReg, the pipeline generates genome-wide input in ∼6 Mb “chunks”, using the central ∼2 Mb region for prediction, though chunk widths scale automatically with user-defined resolutions.

**Figure 1:**
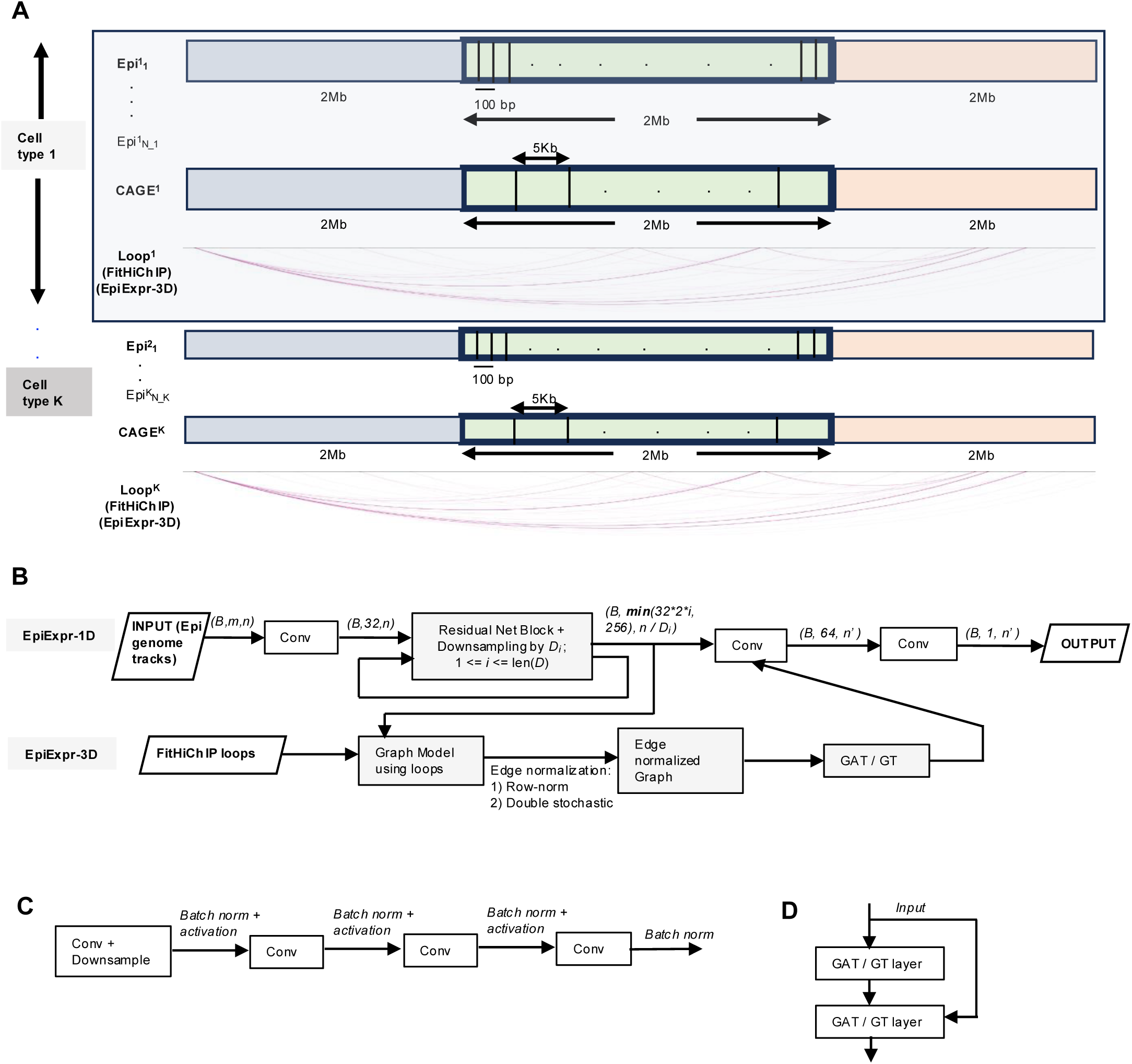
Schematic of the proposed EpiExpr (1D and 3D) models using residual net blocks (CNN) and initial residual connection (GNN) to predict gene expression from epigenetic data. **(A)** EpiExpr supports creating training and validation datasets from multiple cell types, and for each cell type *k*, supports input of variable *N_k* number of epigenetic tracks. Each cell type *k* has at most one CAGE track CAGE*^k^*. EpiExpr-3D additionally requires FitHiChIP loops as an input for each cell type, having the same resolution as the expression track. The dataset supports arbitrary epigenetic and expression track resolutions, but the current study uses 100bp and 5kb, respectively. Similar to Epi-GraphReg, each chunk uses 6Mb for training, and the middle 2Mb segment is used for predicting gene expression. **(B)** Schematic of EpiExpr-1D and 3D prediction models. EpiExpr-1D employs convolution and residual net to downsample and predict gene expression in the target expression resolution. The residual net block iteratively performs convolution and downsampling by a factor *D_i_* at each step where *D* is the vector of downsampling factors and *i* is the iteration counter. EpiExpr-3D additionally uses chromatin looping information right after the residual net block, and supports both GAT and GT models with different graph edge normalization techniques. The triplets (*B,m,n*) refer the data dimension (PyTorch) at any step - *B* refers to batch size (set as 1), *m* is the input channels (depends on the number of input epigenetic tracks), and *n* is the sequence length (depends on the track resolution). (C) Detailed view of the residual net block consisting of multiple layers of convolution and batch normalization. (D) Schematic of initial residual connection in GAT or GT models, where the initial signal is applied on each layer of GAT / GT.

The EpiExpr-1D model architecture is built on a residual CNN framework with iterative, adaptive downsampling to match input resolution *e* to output resolution *c* (**Fig. 1B**). For each (*e, c*) pair, the model computes the prime-factor sequence required to reduce resolution from *e* to *c* and assigns these factors to successive residual blocks (**Methods**). For example, when converting 100 bp inputs to 5 kb outputs, EpiExpr-1D uses three residual blocks with downsampling factors of 2, 5, and 5. A minimum of three residual blocks is enforced; if the required downsampling vector has fewer than three elements, additional blocks with a downsampling factor of 1 are inserted. Each block follows a ResNet18-style architecture^40^ consisting of four convolutional layers (with downsampling applied in the first), along with batch normalization and activation layers (**Fig. 1C**). The complete set of residual net blocks are preceded and followed by 1 and 2 layers of convolution, respectively, to combine the information of *m* epigenetic tracks to a single-channel expression prediction (**Methods**).

We benchmarked EpiExpr-1D on GM12878 and K562 using the epigenomic and expression tracks provided in the reference study (hg19 reference genome)^36^. For comparison, we extended the reference 1D Epi-GraphReg model^36^ to a custom Snakemake pipeline (https://github.com/souryacs/EpiGraphReg_Custom) for curating training data, supplying identical sets of epigenetic (100 bp) and CAGE (5 kb) tracks and training separate models per cell type (**Methods**). Evaluation metrics included Pearson correlation and mean absolute error (MAE) between predicted and observed log2 expression. We assessed expression bins having at least one protein-coding TSS (TSS >= 1). We also considered different subsets of expression bins according to their true expression levels: all bins (Expr >= 0), bins with expression >= 1 and bins with expression >= 5, in order to assess whether EpiExpr can recover expressions of both low and highly expressed bins.

Across both cell types and all bin subsets, EpiExpr-1D consistently achieved higher correlations than Epi-GraphReg-1D, and lower MAE in nearly all conditions, except for GM12878 with highly expressed bins (Expr>= 5) where MAE values were marginally higher than those of Epi-GraphReg-1D (**Figs. 2A–B; Suppl. Table 1**). This comprehensive evaluation demonstrates the effectiveness of the proposed residual-CNN architecture for modeling gene expression solely from 1D epigenomic signals.

**Figure 2:**
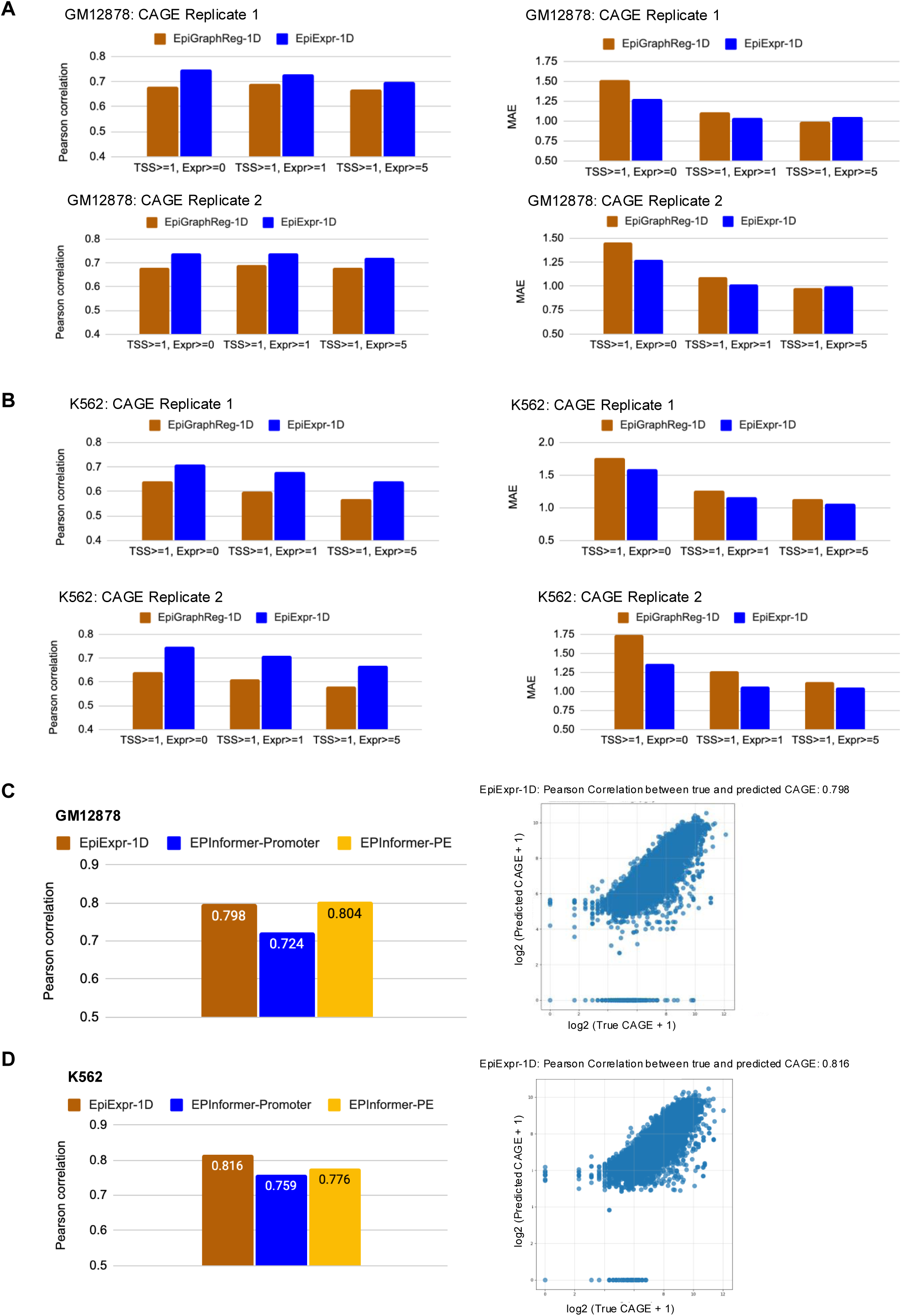
Comparison between EpiExpr-1D, Epi-GraphReg-1D, and EPInformer techniques for GM12878 and K562 datasets obtained from Epi-GraphReg and Basenji papers. **(A)** Comparison of Pearson correlation (left) and mean absolute error (MAE) (right) between the true and predicted expressions for Epi-GraphReg-1D and EpiExpr-1D, with respect to the epigenomic datasets and two CAGE tracks for GM12878 cell type, provided in the Epi-GraphReg paper. Track resolutions: epigenetic 100 bp, CAGE 5Kb. Validation chr: 1, 11. test chr: 2, 12. TSS>= *x,* Expr >= *y*: expression bins with at least *x* number of overlapping TSS and minimum expression *y*. **(B)** Similar comparisons as (A) for K562 cell type specific two CAGE tracks. **(C)** Comparison of mean (across 12-fold cross validation) Pearson correlations between true and predicted expressions (in log2 scale) for EpiExpr-1D and two versions of EPInformer, namely EPInformer-Promoter (using promoter sequences) and EPInformer-PE (using both promoter and enhancer sequences) with respect to the epigenomic and CAGE datasets for GM12878 cell type, provided in the Basenji paper. EPInformer measures correlation of expressions per gene, while EpiExpr models measure correlations per 5 Kb expression bins (left); True and predicted log2 expressions of EpiExpr-1D across the complete 12-fold cross validation, where individual points refer to 5 Kb expression bins (right). **(D)** Similar comparisons as (C) for the K562 cell type specific datasets.

We next compared EpiExpr-1D to the sequence-based EPInformer model^31^, which integrates DNA sequence, epigenomic tracks, and 3D chromatin interactions using dilated convolutions and multi-head attention. Using epigenomic and expression tracks from the Basenji repository^15^ for the cell types GM12878 and K562 (hg38 reference genome), we generated matched training and validation datasets with 100bp epigenetic and 5kb expression track resolutions. Performance was evaluated using 12-fold cross-validation, consistent with prior work (**Methods**), and measured as Pearson correlation between predicted and true expression values for bins overlapping unique protein-coding TSS from Xpresso^41^, matching the evaluation protocol of EPInformer.

We compared EpiExpr-1D against two EPInformer variants: (1) EPInformer-Promoter, which uses promoter-proximal DNA sequence, and (2) EPInformer-PE, which incorporates promoter and enhancer sequences along with promoter-enhancer distances. Using the performance values reported in the EPInformer manuscript^31^, we observed that EpiExpr-1D achieves similar correlation to both EPInformer variants in GM12878 and higher correlation in K562 (**Figs. 2C-D; Suppl. Table 2**). Importantly, EPInformer requires transformer layers over long DNA sequences and an additional preprocessing step using the ABC model^33,34^ to define promoter-enhancer candidates, making it computationally intensive. By contrast, EpiExpr-1D relies solely on epigenomic profiles and a lightweight residual CNN architecture, reducing computational cost while achieving comparable or superior predictive performance. Thus, EpiExpr-1D provides an efficient and practical alternative for gene-expression prediction from 1D epigenetic data.

### EpiExpr-3D: integrating 3D chromatin interactions to improve gene expression prediction

We next developed EpiExpr-3D to evaluate whether incorporating chromatin interactions improves epigenome-based prediction of expression. Prior approaches such as Epi-GraphReg used graph attention networks (GAT) applied to HiCDC+^42^ interactions derived from HiChIP or Hi-C datasets. Here, we instead leveraged loop calls from our previously published tool FitHiChIP^43^, which supports multiple 3C assays (Hi-C, PCHi-C, HiChIP) and reports contact counts, statistical significance, and the bias values of interacting segments, which all can be used to define the edge features for the graph (next section). FitHiChIP has been shown to achieve superior loop-calling performance across diverse benchmarking settings^43^, providing a robust basis for defining edges in our chromatin-interaction graphs. In EpiExpr-3D, the intermediate representations produced by the EpiExpr-1D residual block at CAGE resolution serve as node embeddings for a graph neural network (GNN), whose output is passed to the later convolutional layers of EpiExpr-1D to generate final predictions (**Fig. 1B**). This end-to-end training strategy avoids the instability observed when applying a separately pre-trained CNN (such as pre-trained EpiExpr-1D model) on top of a GNN, including zero-gradient collapse and NaN outputs.

EpiExpr-3D supports two attention-based GNN architectures: a graph attention network (GATv2Conv)^38^ and a graph transformer (TransformerConv)^39^ implementing a unified message passing model combining both GNN and label propagation algorithm (LPA). We employed eight attention heads and two GNN layers for both models. Graph edges were constructed from significant (FDR < 0.1) H3K27ac HiChIP loops in GM12878 and K562 (**Methods**). We compared two edge-normalization approaches - row-normalization from scikit-learn (referred as E1) and double-stochastic normalization (denoted by E2)^44^ (**Methods**). We also optionally incorporated an initial residual connection (referred as R) that inputs the initial features at each GNN layer (**Fig. 1D**) which was previously shown to improve GNN performance^45^.

We first benchmarked the utility of EpiExpr-3D and corresponding GNN architectures using the epigenetic and expression datasets provided in the Epi-GraphReg^36^ study and applying on our custom Snakemake pipeline (https://github.com/souryacs/3CExpr/tree/main/EpiExpr_3D_Create_Training_Data) to create training and validation datasets. Results for different settings of GAT revealed that GAT with residual connections (R) involving either of the edge normalization techniques (E1 or E2) performed better than GAT without residual connection (**Suppl. Fig. 1**). Specifically, GAT with scikit-learn edge normalization and residual connection (E1+R) consistently produced higher correlations and lower MAE across both datasets while the GAT model with double stochastic normalization (E2+R) showed marginally higher performance for K562 datasets (**Suppl. Fig. 1**). Thus, for the GAT model, we have finalized the E1+R setting for the subsequent results and benchmarking.

Performances of different settings for the graph transformer (GT) model, on the other hand, showed greater variability, but overall for both GM12878 and K562 datasets, results for the settings E1 (scikit-learn edge normalization without residual connection) and E2+R (double stochastic edge normalization with residual connection) of GT architecture showed superior performance in terms of either higher correlation with expression or lower MAE (**Suppl. Fig. 2**). Thus, we tested both E1 and E2+R settings for the GT model in the subsequent results.

We also evaluated whether incorporating loop-level attributes such as contact counts, statistical significance, etc. as edge features in the GNN would enhance prediction accuracy. Specifically, we tested the following loop-derived quantities: (1) p-value of interaction significance, (2) observed-over-expected contact counts, and (3) bias-normalized contact counts (**Methods**). Although the combination of contact-count–based features (2 and 3) yielded the strongest performance among the feature sets for the GAT architecture, none of these loop derived features when applied on the GAT model (denoted by the setting **+F**) consistently outperformed the respective GAT models without any edge features, in either GM12878 or K562 datasets from Epi-GraphReg (**Suppl. Fig. 3**). Similar trends were observed for the graph transformer (GT) models as well (**Suppl. Fig. 4**). Consequently, subsequent analyses used EpiExpr-3D without incorporating additional edge features.

Results using Epi-GraphReg epigenomic datasets^36^ showed that integrating chromatin interactions via GNNs in EpiExpr-3D marginally improved expression prediction relative to EpiExpr-1D for K562, particularly when using the graph transformer architecture, and for non-expressing CAGE bins (**Figs. 3A-B**). For expressed (Expr >= 1) bins, EpiExpr-1D in fact, showed higher correlation and lower or similar MAE compared to EpiExpr-3D + GT models (**Figs. 3A-B**). For GM12878 data, EpiExpr-1D produced comparable or higher correlation and lower MAE than all EpiExpr-3D settings (**Suppl. Fig. 5**). These differences suggest that chromatin interactions may be informative of the expression status (zero vs non-zero) of bins. Importantly, both EpiExpr-1D and EpiExpr-3D surpassed the 1D and 3D Epi-GraphReg models across both cell types (**Figs. 3A-B, Suppl. Fig. 5**), underscoring the advantage of the residual network backbone over conventional CNN-plus-GAT designs.

**Figure 3:**
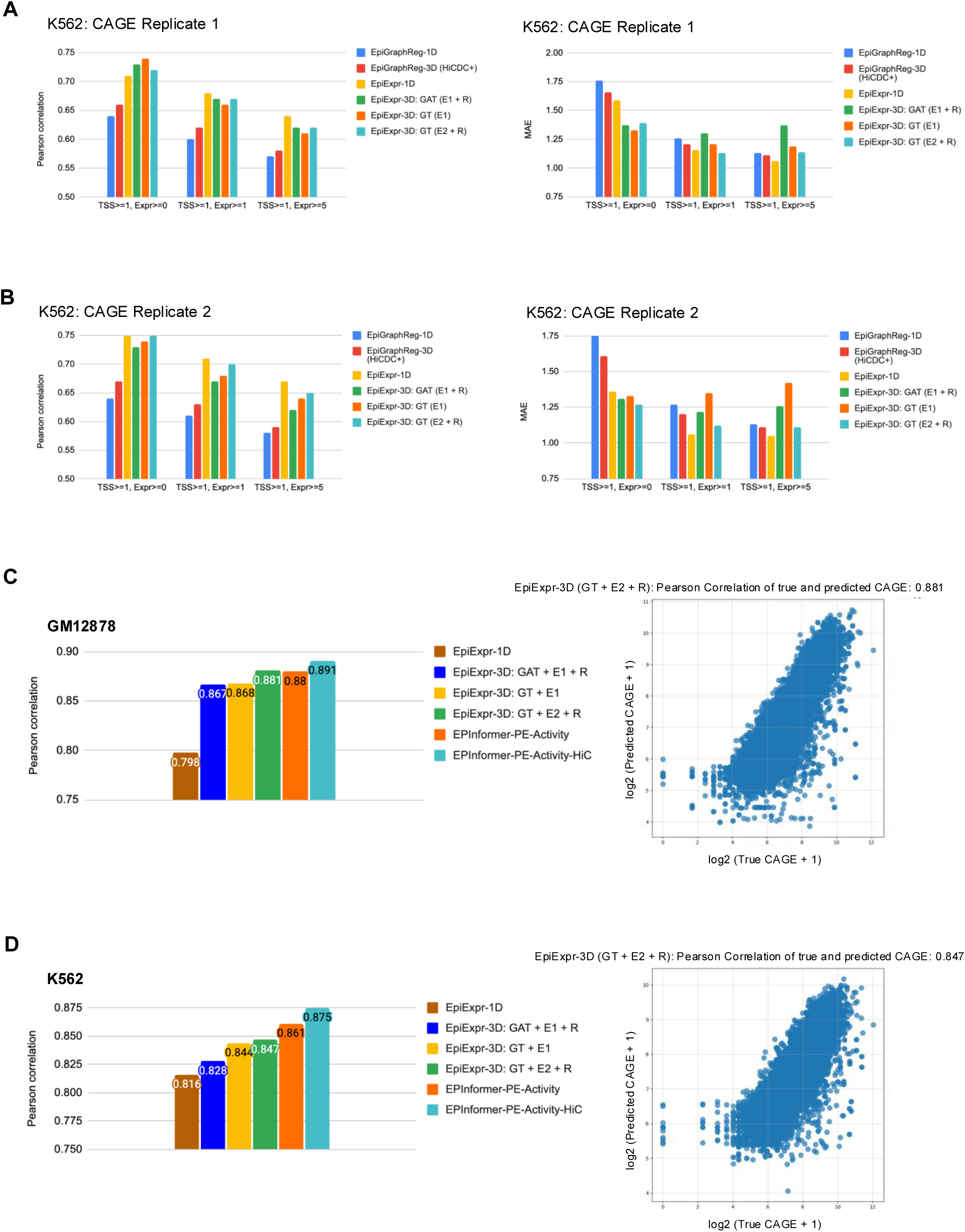
Comparison between EpiExpr-3D, Epi-GraphReg-3D, and EPInformer (+ ABC Score) techniques for GM12878 and K562 datasets obtained from Epi-GraphReg and Basenji papers. **(A)** Comparison of Pearson correlation (left) and mean absolute error (MAE) (right) between the true and predicted expressions for Epi-GraphReg (1D and 3D with HiCDC+ loops) and different settings of EpiExpr-3D, with respect to the epigenomic datasets and CAGE replicate 1 for K562 cell type, provided in the Epi-GraphReg paper. Track resolutions: epigenetic 100 bp, CAGE 5Kb. Validation chr: 1, 11; test chr: 2, 12. TSS>= *x,* Expr >= *y*: expression bins with at least *x* number of overlapping TSS and minimum expression *y*. GAT: graph attention network, GT: graph transformer, E1: scikit-learn row normalization of graph edges, E2: double stochastic normalization of graph edges, R: using initial residual connection on GAT / GT. **(B)** Similar to (A) for the second CAGE replicate of K562 cell type. **(C)** Comparison of mean (across 12-fold cross validation) Pearson correlations between true and predicted expressions (in log2 scale) for various settings of EpiExpr-3D and two versions of EPInformer, namely EPInformer-PE-Activity (using both promoter and enhancer sequences and corresponding regulatory activities coupled with ABC score) and EPInformer-PE-Activity-HiC (also using the HiC contacts and ABC scores between the promoters and enhancers) with respect to the epigenomic and CAGE datasets for GM12878 cell type, provided in the Basenji paper. EPInformer measures correlation of expressions per gene, while EpiExpr models measure correlations per 5 Kb expression bins (left); True and predicted log2 expressions of EpiExpr-3D: GT + E1 across the complete 12-fold cross validation, where individual points refer to 5 Kb expression bins (right). **(D)** Similar comparisons as (C) for the K562 cell type specific datasets.

We next benchmarked EpiExpr-3D on the Basenji epigenomic tracks^15^ and compared its performance with two EPInformer variants that incorporate ABC-derived promoter and enhancer activity scores^33^, without (denoted by EPInformer-PE-Activity) or with (denoted by EPInformer-PE-Activity-HiC) Hi-C signal. Unlike these methods which rely on pre-computed sequence embeddings and ABC-generated activity and contact features, EpiExpr-3D is trained solely on epigenomic signals and chromatin-interaction graphs, without using sequence-based transformers or external feature-generating models. Across GM12878 and K562 with 12-fold cross validation, adding graph-based information in either GAT or GT settings considerably improved performance over EpiExpr-1D (**Figs. 3C-D**), with the graph transformer (GT) using double-stochastic normalization (E2) and residual connections (R) achieving the highest accuracy among our GNN settings (**Figs. 3C-D**). EpiExpr-3D matched the performance of EPInformer for GM12878 (**Fig. 3C**) and approached it for K562 (**Fig. 3D**), despite its substantially lower computational complexity and lack of dependence on ABC-derived features. These results highlight that a compact residual CNN plus GNN architecture operating directly on epigenomic and interaction data provides an efficient and competitive solution for gene expression prediction without requiring DNA sequence embeddings and computationally intensive transformer architecture.

### EpiExpr-1D and EpiExpr-3D retrieve CRISPRi validated regulatory enhancers

To evaluate whether EpiExpr-1D and EpiExpr-3D recover experimentally validated interactions between genes and enhancers (E) or promoters (P), we benchmarked both models against reference CRISPRi-FlowFISH enhancer perturbation screens from the Fulco et. al. study^33^. These assays use KRAB-dCas9-mediated interference of candidate enhancer regions followed by RNA FISH to quantify their perturbation impact, and also report activity-by-contact (ABC) scores per E-G pair based on the geometric mean of H3K27ac and DNase signals in promoter and enhancer regions and KR-normalized Hi-C contact frequency between them. Similar to the reference study, we considered a regulatory element (enhancer) significant if it is mentioned as *significant* in the input data and downregulates the target gene expression. Using genes having at least 10 validated enhancers and one statistically significant candidate enhancer downregulating target gene expression, we curated two different lists for validation. The first list contains 3,995 pairs corresponding to 25 genes and regulatory elements (E or P) while the second list excludes all the promoters (P) and considers 2,671 E-G pairs corresponding to 21 genes for validation (**Methods**). In contrast to sequence-based approaches such as Enformer and EPInformer, we did not restrict enhancer candidates by distance to the transcription start site. For both EpiExpr-1D and EpiExpr-3D models, we computed DeepSHAP^23^ attribution scores at 100-bp resolution using an all-zero baseline to assess the contribution of each epigenomic peak (enhancer) to predicted expression. As both EpiExpr-1D and EpiExpr-3D models report gene expression in terms of 5-Kb bins, similar to the Epi-GraphReg study, we used the AUPRC script of Epi-GraphReg (**Methods**) to benchmark the methods against the reference ABC scores^33^.

Considering the reference 3,995 E/P-G pairs, EpiExpr-1D performed marginally better than the ABC model in identifying functional regulatory regions, achieving higher mean AUPRC values (0.3677 vs. 0.3508; **Fig. 4A**) and much higher gene-wise median AUPRC (0.5446 vs. 0.3495; **Fig. 4B**). EpiExpr-3D with GAT architecture yielded slightly lower mean AUPRC (0.2847 for GAT+E1+R) but high gene-wise median AUPRC (0.5087) comparable with EpiExpr-1D (**Fig. 4A**). This modest reduction of mean AUPRC is attributable to two factors: (1) message-passing operations in GAT produce more diffuse attribution patterns, reducing enhancer specificity, (2) attributions are computed at 100-bp epigenomic resolution, whereas the GNN operates at 5-kb resolution, making cross-resolution mapping less precise. We also evaluated graph transformer (GT) models, which adapt transformer-style attention for GNNs. GT architectures improved prioritization of regulatory elements compared to GAT, yielding higher mean AUPRC (0.3582 for GT+E1 and 0.343 for GT+E2+R; **Fig. 4A**) and comparable gene-wise median AUPRC (0.4502 for GT+E1 and 0.4712 for GT+E2+R; **Fig. 4B**) in addition to their previously observed gains in gene-expression prediction (**Figs. 3C-D**).

**Figure 4:**
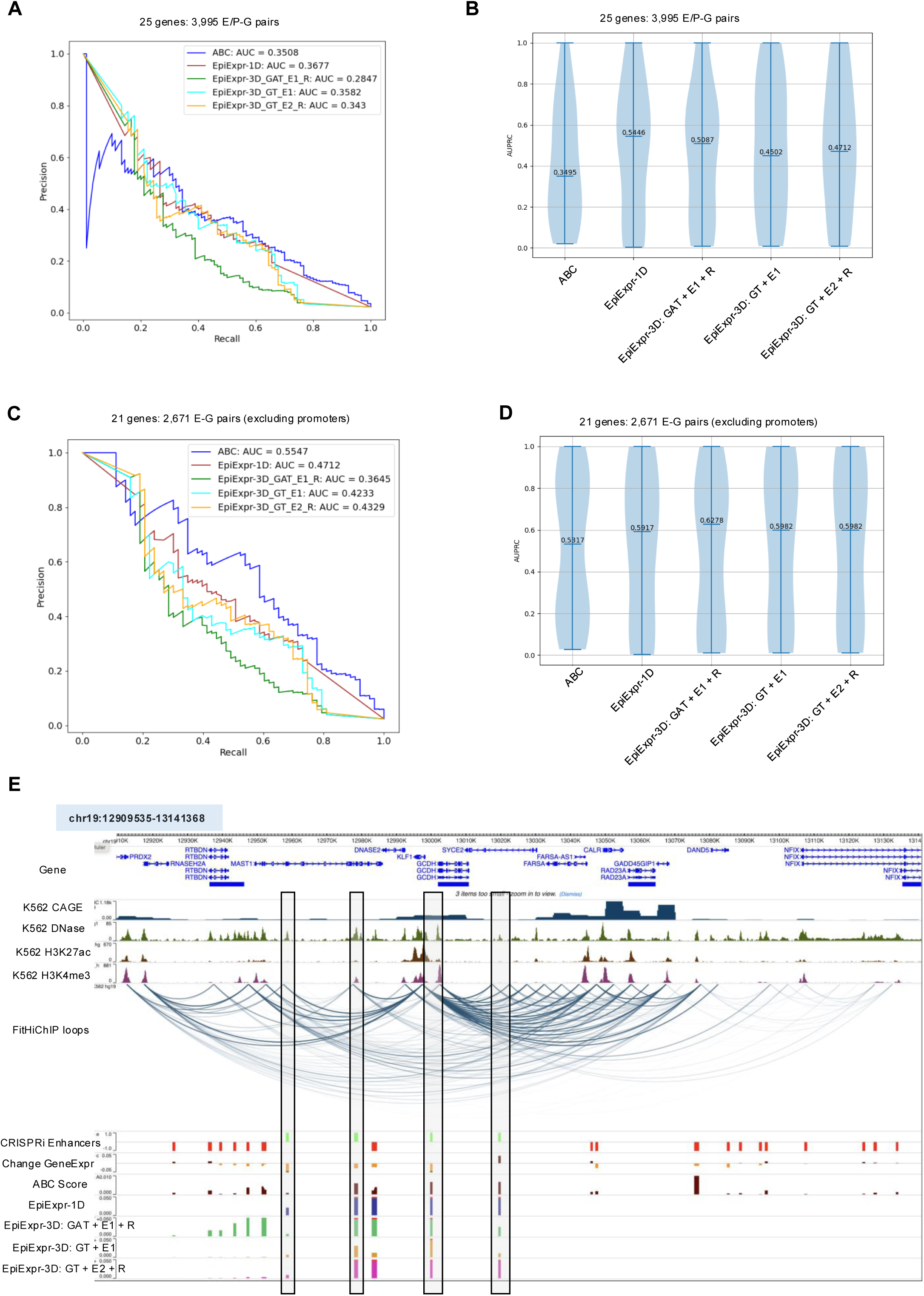
Benchmarking EpiExpr-1D, EpiExpr-3D, and ABC models by AUPRC and locus specific examples, with respect to the CRISPRi validated enhancers from K562 cell type: **(A)** Precision-Recall plot and area under curve (AUPRC) between Activity-by-Contact (ABC) Scores and the proposed EpiExpr-1D and various settings of EpiExpr-3D models, with respect to CRISPRi validated 3,995 E/P-G pairs in^33^. **(B)** Boxplot of median AUPRC values per gene, comparing between ABC, EpiExpr-1D, and various settings of EpiExpr-3D models, with respect to the data in (A). **(C)** Similar analysis as (A) but with respect to the CRISPRi validated 2,671 E-G pairs (excluding the promoters). **(D)** Similar plot as (B), with respect to the data in (C). **(E)** WashU browser track from *KLF1* locus and its up and downstream 200 Kb, showing the input K562 expression and epigenetic tracks, FitHiChIP loops, CRISPRi validated enhancers (green lines indicate the significant ones while the red lines denote the non-significant entries), change of gene expression as per corresponding enhancer perturbation, ABC scores from^33^, DeepSHAP attribution scores from EpiExpr-1D and EpiExpr-3D models in 100 bp resolution. Boxed regions highlight the CRISPRi validated significant enhancers.

When restricting the analysis to the 2,671 reference enhancer–gene (E-G) pairs excluding promoters, the ABC model achieved higher mean AUPRC (0.5547) compared to all EpiExpr-1D and EpiExpr-3D configurations (**Fig. 4C**). EpiExpr models (1D or 3D), however, produced marginally higher gene-wise median AUPRC compared to the ABC model (**Fig. 4D**), indicating overall high average recovery of regulatory elements per gene for different EpiExpr models. We note that mean AUPRC values for EpiExpr-3D (GAT or GT) settings are marginally lower than EpiExpr-1D and ABC models. Although graph-based models (GAT and GT) are designed to capture long-range regulatory interactions, their performance is influenced by above-mentioned factors of message passing and cross-resolution mapping. Further, ABC scores are computed on precisely annotated enhancer and gene intervals, whereas DeepSHAP attributions computed on 100bp resolution do not explicitly model enhancer boundaries, an issue that is exacerbated for distal regulatory elements. Direct comparison with EPInformer^31^ was not feasible as it evaluated only 774 E-G pairs and did not report the full gene list. Epi-GraphReg, on the other hand, considered 2,574 E-G pairs (a subset of the tested 2,671 E-G pairs in this study) for evaluation. In spite of following the validation script from Epi-GraphReg, we observed discrepancies in ABC performance: our analysis yielded an AUPRC of 0.5547 (**Fig. 4C**), compared with 0.364 reported in Epi-GraphReg (**Methods**). Notably, EpiExpr-1D achieved higher mean AUPRC (0.4712) than Epi-GraphReg (0.4185) as reported previously^36^.

We next examined the performance of EpiExpr and the reference ABC model at the *KLF1* locus within a 250 kb genomic window. Both ABC and multiple EpiExpr configurations successfully identified validated regulatory enhancers proximal to, and approximately 50 kb upstream of, the *KLF1* promoter. In contrast, the ABC model additionally predicted regulatory elements located 70-100 kb downstream of the promoter that are not supported by experimental evidence, whereas these false positives were not detected by EpiExpr, indicating improved specificity (**Fig. 4E**). Despite the known limitations of DeepSHAP-based attribution arising from cross-resolution mapping, these results underscore the ability of EpiExpr models to recover distal regulatory enhancers while maintaining higher precision.

We note that EPInformer computed ABC scores using the hg38 reference genome and applied distinct thresholding and aggregation strategies for benchmarking. Owing to limited computational resources, particularly GPU memory constraints, we were unable to compute DeepSHAP attribution scores for K562 trained models from Basenji data (hg38) with hundreds of epigenetic tracks. Consequently, we relied on the K562 trained models from Epi-GraphReg data (hg19) and adopted the benchmarking protocol described in the Epi-GraphReg manuscript for performance comparison (**Methods**).

### EpiExpr-1D and EpiExpr-3D incur very low computational resources and running times

Curation of the training data for EpiExpr-3D using the Snakemake pipeline on the Epi-GraphReg datasets required less than 30 minutes and approximately 1 GB of CPU memory. Inference with the EpiExpr-3D model under the GAT/GT configuration completed in ∼40 minutes, using ∼1 GB of CPU memory and a single GPU with a peak memory allocation of 10 GB. Despite these GPU memory constraints, EpiExpr models offer an efficient and computationally lightweight framework for accurate gene expression prediction.

## DISCUSSION

Recent transformer-based approaches have demonstrated the ability to predict gene expression from one-hot–encoded DNA sequences and fixed-resolution epigenetic tracks (e.g., 128 bp)^27^. However, these architectures demand substantial computational resources and large, multi-cell-type training datasets with hundreds of epigenetic tracks. In contrast, recent studies suggest that supervised CNN models can achieve comparable performances to pre-trained, unsupervised transformers^46^. Motivated by this, we developed EpiExpr (1D and 3D) using residual CNNs as the backbone, applied exclusively to epigenetic tracks without DNA sequence embeddings, to directly evaluate the power of epigenomic information and chromatin interactions in predicting gene expression.

While Epi-GraphReg^36^ previously explored gene expression prediction from epigenetic tracks and 3D interactions, it was constrained by a fixed set of cell types and track resolutions. Our open-source Snakemake pipelines for curating datasets address these limitations, enabling flexible training datasets across multiple cell types, arbitrary numbers of epigenetic tracks, and user-defined epigenetic and CAGE resolutions. Both EpiExpr-1D and EpiExpr-3D consistently outperformed Epi-GraphReg across multiple datasets. We further benchmarked EpiExpr-3D against EPInformer, which integrates DNA sequence embeddings, epigenetic features, chromatin interactions, and ABC-derived activity scores. Due to computational constraints, we relied on published performance metrics rather than running EPInformer directly, and followed their performance metrics and evaluation guidelines (such as 12-fold cross validation) while running EpiExpr-1D and 3D models. Remarkably, EpiExpr-3D employing the graph transformer (GT) architecture matched EPInformer-PE-Activity-HiC performance while requiring substantially lower computational resources, highlighting its applicability for large-scale, multi-cell-type applications.

For interpretability, we primarily used DeepSHAP^23^ for attribution. We also tested Integrated Gradients^47^ and observed similar AUPRC but with marginally higher sparsity (zero attribution scores). In addition to using the attention scores from the GAT or GT models (5kb resolution), we also attempted to employ GNNExplainer^48^ but observed that it is not suitable for EpiExpr-3D due to the interleaving of CNN and GAT (or GT) layers while GNNExplainer expects only the GNN model for gradient backpropagation.

This study also lays the foundation for several future directions. First, transformer architectures could be applied directly to the EpiExpr-1D datasets to benchmark against CNNs, potentially leveraging FlashAttention^49^, but would require substantially higher computational resources. Second, higher-resolution epigenetic and chromatin interaction datasets (such as 100bp epigenetic and 500bp CAGE resolutions) could further refine predictions, contingent on computational resources and the availability of high-resolution chromatin interaction datasets. Third, multi-cell-type training and evaluation remain unexplored; future work will extend EpiExpr to such datasets to assess generalizability, similar to the study in^50^. Fourth, applying GNN models at the epigenetic resolution before downsampling to expression resolution may provide deeper insights into the relative merits of graph attention or graph transformers versus sequence-based transformers, although this would require substantially higher computational capacity.

Overall, EpiExpr-1D and EpiExpr-3D, together with the accompanying data-curation pipelines, constitute a flexible and scalable framework for predicting gene expression from epigenetic and chromatin interaction data. These models support multiple cell types, arbitrary numbers of tracks, and user-defined resolutions, and they can be further extended with advanced transformer or graph-transformer architectures as computational resources allow.

## METHODS

### Datasets

*Blacklist files for various reference genomes:* Blacklist files for the reference genomes hg19 and hg38 were downloaded from the Zenodo repository https://zenodo.org/records/1491733. These regions were excluded from the generated epigenetic track embeddings.

*GM12878 and K562 epigenomic dataset (hg19) from Epi-GraphReg study:* We downloaded the datasets for GM12878 and K562 cell types (aligned to hg19), following the supplementary material of Epi-GraphReg paper^36^. We downloaded the 1D epigenomic tracks (DNase-seq, ChIP-seq for histone marks H3K27ac, H3K4me3, and transcription factor JUND) in bigWig format. For GM12878, we downloaded two CAGE files: ENCFF915EIJ (replicate 1) and ENCFF990KLZ (replicate 2). For K562, we downloaded two CAGE files: ENCFF623BZZ (replicate 1) and ENCFF902UHF (replicate 2). Expression bam files were normalized using the *bamCoverage* routine from DeepTools^51^ to account for sequencing depth (reads per genome coverage, RPGC). Epigenomic and CAGE tracks were processed at 100 bp and 5 kb (5000 bp) resolutions, respectively.

*HiChIP interactions for GM12878 and K562 (hg19):* H3K27ac HiChIP valid pairs for individual replicates of GM12878 and K562 were obtained from the reference HiChIP study^8^. For GM12878, two replicates were merged, while three replicates were merged for K562. Merged valid pairs were processed with our previously published HiChIP loop caller FitHiChIP (v9.1)^43^. ChIP-seq input files were the same as listed in Supplementary Table 1 of FitHiChIP manuscript^43^. HiChIP interactions were reported at 5 kb resolution (BINSIZE=5000) using peak-to-all output (IntType=3), distance range 20 kb - 2 Mb, loose background (UseP2PBackgrnd=0), coverage bias regression (BiasType=1), no merge filtering (MergeInt=0), and 10% FDR (QVALUE=0.1).

*GM12878 and K562 epigenomic dataset (hg38) from the Basenji paper:* For hg38, epigenomic and expression tracks for GM12878 and K562 were downloaded from the Basenji^15^ repository: https://raw.githubusercontent.com/calico/basenji/0.5/manuscripts/cross2020/targets_human.txt. We downloaded CAGE files with accession numbers CNhs12333 for GM12878 and CNhs11250 for K562 cell type.

*HiChIP interactions for GM12878 and K562 (hg38):* H3K27ac HiChIP valid pairs for GM12878 and K562 aligned to hg38 were obtained from our previously published Loop Catalog repository^52^. These valid pairs were processed with FitHiChIP using the same settings as described above.

### EpiExpr-1D model training data and Snakemake pipeline

We developed a Snakemake pipeline to generate training datasets for the EpiExpr-1D model, and the pipeline has been made available as an open-source repository (https://github.com/souryacs/3CExpr/tree/main/EpiExpr_1D_Create_Training_Data). The pipeline requires the following input parameters: (1) reference genome (hg19 or hg38), (2) chromosome sizes and transcription start site (TSS) information for the chosen genome, (3) resolutions for epigenomic tracks (e.g., 100 bp) and CAGE tracks (e.g., 5 kb), and (4) file paths for the epigenomic and expression tracks corresponding to the specified resolutions.

Following Epi-GraphReg^36^, the pipeline segments each chromosome into 6 Mb windows with a 2 Mb step size, corresponding to epigenetic track resolution of 100bp and CAGE resolution of 5kb. The central 2 Mb region of each window is used for gene expression prediction, while the flanking 2 Mb regions on each side serve as background. For example, at 100 bp and 5 kb resolutions for epigenomic and CAGE tracks, each 6 Mb window yields an epigenomic vector of length 60,000 and an expression vector of length 1,200. The pipeline supports any cell type, any number of epigenomic tracks, and flexible track resolutions. Data processing and serialization are performed using PyTorch (version 2.4.1) routines, with each chromosome and data chunk saved in .pkl format.

### EpiExpr-3D model training data and Snakemake pipeline

Similar to EpiExpr-1D, we developed a Snakemake pipeline to generate training datasets for the EpiExpr-3D model, which has been deployed as an open-source repository (https://github.com/souryacs/3CExpr/tree/main/EpiExpr_3D_Create_Training_Data). In addition to the parameters required for EpiExpr-1D, this pipeline incorporates chromatin interactions for the respective cell types. For this study, we used HiChIP interactions derived from FitHiChIP with an FDR threshold of 0.1, although the pipeline is compatible with interactions from any 3C-based protocol, including Hi-C and PCHi-C. Chromatin interactions are required to match the expression resolution. We also provide a custom script to convert FitHiChIP output interactions into the specific BED format used by this pipeline.

For the central 2 Mb segment of a 6 Mb window (used for gene expression prediction), chromatin interactions within 2 Mb upstream and downstream are incorporated to model the graph neural network. Training data for each chromosome, including both epigenetic tracks and chromatin interactions, is saved in PyTorch .pkl format for each window segment.

### Adaptation of Epi-GraphReg model

The Epi-GraphReg^36^ model predicts gene expression using epigenomic data but the current implementation only supports hg19 reference genome and a fixed set of six epigenomic tracks per cell type (GM12878 and K562). We adapted the Epi-GraphReg package into a new open-source repository (https://github.com/souryacs/EpiGraphReg_Custom) that supports any number of epigenetic tracks, multiple cell types, user-defined epigenetic and CAGE track resolutions, and outputs predicted expression at the target resolution. Two versions were created for direct comparison with EpiExpr: Epi-GraphReg-1D, which uses only 1D epigenetic data, and Epi-GraphReg-3D, which additionally incorporates chromatin interactions. For Epi-GraphReg-3D, we used HiCDC+^42^ interactions as per their default implementation and recommendation by the authors, on GM12878 and K562 HiChIP datasets with a 0.1 FDR threshold and distance range of 20 kb - 2 Mb, following Bioconductor documentation.

### Performance comparison with Epi-GraphReg

To benchmark EpiExpr (1D and 3D) against Epi-GraphReg, we employed Pearson correlation and mean absolute error (MAE) between true and predicted expression (binned, log2 scale). We also computed these metrics for subsets of expression bins based on different thresholds of expression (≥0, ≥1, ≥5). Each bin has at least one TSS (TSS >= 1). Expression bins overlapping blacklisted regions (hg19) were excluded.

### EpiExpr-1D model architecture

EpiExpr-1D processes the 6 Mb training segments (PyTorch .pkl files) sequentially (batch size = 1). To account for different numbers of epigenetic tracks for different datasets, and also to support using training data from different cell types or conditions (having different numbers of epigenetic tracks in the respective datasets), the model first projects input tracks into *M* channels, where *M* is the smallest power of 2 greater than the number of input tracks (for one cell type) or the maximum number of input tracks across different cell types. For Epi-GraphReg GM12878 and K562 (hg19) data, we employed *M* = 8; for Basenji GM12878 and K562 (hg38) datasets, we used *M* = 256 and 512, respectively, according to the number of epigenetic tracks in the respective cell types.

The projected data is then processed through a residual network. We used PyTorch routine *Conv1d* to perform the convolution. Convolutions are performed with kernel size = 5, GELU activation, dropout = 0, learning rate = 1e-3, L2 regularization = 1e-5, and Huber loss as the loss function, unless otherwise specified. The model begins with a convolution layer (32 output channels), followed by batch normalization and activation. Residual net blocks perform adaptive downsampling to convert from epigenomic resolution *e* to CAGE resolution *c*. For instance, if *e* = 100bp and *c* = 5000bp, downsampling factors of 2, 5, and 5 are applied sequentially. EpiExpr-1D applies at least three layers of residual net block. So if the length of the downsampling vector is less than 3, additional residual net layers with downsampling factor of 1 are utilized. Each residual block doubles the input channels, with a maximum of 256. Following the architecture of the ResNet18 model, each block consists of four convolution layers, four batch normalization layers, and three activation layers, with the first convolution performing downsampling, by setting the *stride* parameter of the routine *Conv1d* equal to the downsampling factor. The output from the residual blocks is further processed through a convolution layer with the same input and output channel dimension (256) and kernel size of 1, batch normalization and activation. To combine the information from 256 channels to 1 channel, we then applied two more convolution layers with output dimensions of 64 and 1, respectively, and with a kernel size of 1. Batch normalization was applied only for the first convolution. The first convolution used GELU while the second used ELU as activation functions. The output is now compatible with the target expression resolution. Note that we did not perform exhaustive parameter tuning (dropout rate, learning rate, L2 regularization), etc. since our primary focus was on establishing the framework.

### EpiExpr-3D model architecture

EpiExpr-3D first applies the EpiExpr-1D model through the residual blocks, omitting the final two convolution layers, yielding 256-dimensional embeddings at the target expression resolution. Chromatin looping information having the same resolution as the expression and embedded in the training data, defines the graph structure, with EpiExpr-1D outputs as node features and chromatin interactions as graph edges. We implemented two GNN architectures:

1. Graph Attention Network (GAT) implemented using the PyTorch routine *GATv2Conv* with 8 heads and 2 layers, both with and without residual connections. The output dimension (*out_channels*) was kept the same as the input dimension (*in_channels*). For GAT without residual connection, the *concat* parameter was set as true; otherwise it was false. Each GAT layer output was applied a layer normalization followed by GELU activation. For GAT with residual connection, the EpiExpr-1D output was added as the input of the second GAT layer.
2. Graph transformer (GT) combining label propagation with message passing and implemented using the PyTorch routine *TransformerConv*, uses 2 layers, 8 heads, and residual connections as needed. Similar to the GAT model, we set the parameter *concat* according to the use of residual connection, and assigned the parameter *beta* to False. We did not, however, employ any layer normalization or activation function on the output of GT layers.

For either of these GNN models, EpiExpr-3D supports two different graph edge normalization techniques for modeling the chromatin interactions into graph architectures. The first technique is row normalization implemented in scikit-learn where each graph edge 𝐸𝑖𝑗 (*i* and *j* denote node indices) is normalized as:

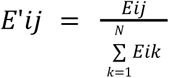

The second technique is double stochastic normalization where each graph edge 𝐸𝑖𝑗 is normalized as:

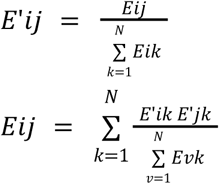

We also tested if adding features from chromatin interactions (such as contact counts) on the graph edges improves the prediction performance or not. So, we tested incorporating various combinations among three chromatin-loop derived features on graph edges: 1) observed contact count / expected contact count for a loop, where the expected contact count is obtained from FitHiChIP output statistics. 2) bias normalized contact count for a loop, defined as: observed contact count / (bias of interacting bin 1 X bias of interacting bin 2), and 3) p-value (statistical significance) of the chromatin loop.

The outputs of GAT or GT have the same resolution as the expression tracks, but have channels of dimension either (256 X 8) or 256, depending on the use of residual connection and corresponding use of the *concat* parameter. So, after the GAT or GT layers, EpiExpr-3D applies two downstream convolution layers of EpiExpr-1D with output dimensions of 64 and 1, respectively, to derive the outputs compatible with the target expression. Both GAT and GT models employ dropout rate = 0, learning rate = 1e-3, Huber loss function between true and predicted outputs, and L2 regularization parameter = 1e-2, which is higher compared to the EpiExpr-1D model due to faster convergence and better performance. However, we did not perform any exhaustive parameter tuning.

### Setting validation and test chromosomes

For comparison with Epi-GraphReg, we employed chromosomes 1 and 11 for validation and chromosomes 2 and 12 for testing, and remaining chromosomes for training. To compare with EPInformer, we employed 12-fold cross-chromosome validation, slightly different from the reference studies^31,36^. For fold 1 to 11, chromosomes *i* and *i* + 10 are used for validation, and chromosomes *i* + 1 and *i* + 11 are set aside for testing. For fold 12, chromosomes 2 and 22 are used for validation, and chromosomes 1 and X are used for testing. Remaining chromosomes are used for training in each fold. Chromosome Y is excluded from all analyses.

### Performance benchmarking with EPInformer

We used the results of the EPInformer method for GM12878 and K562 datasets (hg38 reference genome) as provided in their manuscript^31^. The 12-fold cross-chromosome validation employed in EPInformer is slightly different from our approach since EPInformer also employed chrY for validation. Further, EPInformer reported correlation per gene (TSS), while our proposed EpiExpr models report correlation at the expression bin (5 Kb) level. In spite of these differences, we used the EPInformer results directly from the manuscript, since our limited computational resources did not allow us to re-run EPInformer and/or map the results to expression bins.

### Enhancer prioritization using CRISPRi-FlowFISH data

We obtained the pairs of gene (G) and regulatory enhancers or promoters (E or P) from the CRISPRi-FlowFISH data provided in Suppl. Table 6a of the reference study^33^, listing the regulatory elements altering gene expression in K562 cell type upon perturbation along with the corresponding ABC scores between these pairs. The file contains a total of 5,091 tested pairs of gene and regulatory elements, out of which the true positive (significant + downregulating) regulatory elements are indicated by “TRUE” entries in the “Significant” column, and negative entries in the “fraction of changes in gene expression” of this list. We did not filter these pairs by any distance criterion between the TSS and the enhancers, unlike the reference approaches^27,31^ which only considers enhancers within 100Kb from respective TSS. Rather, we followed the Epi-GraphReg approach and considered only those genes for validation which have at least 10 validated and at least one significant regulatory element. Specifically, we curated two different lists for validation. The first list contained all pairs of genes and regulatory elements satisfying the above criteria, and had 25 genes with 3,995 E/P-G pairs. The second list filtered all promoters from the set of regulatory elements (excluding entries with the field class as “Promoter”) and considered the remaining 2,671 E-G pairs from 21 genes containing the E-G pairs tested in Epi-GraphReg.

To pinpoint the regulatory enhancers corresponding to a gene, we employed the DeepSHAP^23^ approach using the python package *shap* for both EpiExpr-1D and EpiExpr-3D models. For each E-G entry of a particular chromosome *c*, DeepSHAP was applied on the pre-trained model which employed *c* as its test chromosome. For each E/P-G pair, a surrounding 6 Mb chunk (equal to the training / validation data segment length) was used as the input to DeepSHAP, along with zero baseline, as suggested in Epi-GraphReg. Output of DeepSHAP was a vector of attribution scores corresponding to the epigenomic track resolution (100bp). We compared AUPRC trends between ABC score and the proposed EpiExpr models by adapting the benchmarking script provided in the Epi-GraphReg GitHub repository (https://github.com/karbalayghareh/GraphReg/blob/master/feature_attribution/Epi_models_fa_ensemble_FlowFISH.py). Overall attention scores from DeepSHAP in epigenetic track resolution (100bp) were then plotted along with the reference CAGE and epigenetic tracks and FitHiChIP loops using WashU epigenome browser (https://epigenomegateway.wustl.edu/).

Note that EPInformer uses input datasets in hg38 format, and employs hg38 specific CRISPRi resource for benchmarking, as provided in https://github.com/EngreitzLab/CRISPR_comparison/tree/main. We could not, however, run DeepSHAP attribution on the trained models for hg38 K562 Basenji data due to limitations of available computational resources, thus used the hg19 based CRISPRi data provided in^33^ and the script provided in Epi-GraphReg paper for comparison.

### Computational resources

We have used a single NVIDIA A75 GPU with 80GB allocated memory to run EpiExpr-1D and EpiExpr-3D methods. We used PyTorch (version 2.4.1) for implementing CNN-based models and PyTorch Geometric (version 2.6.1) for implementing the GNN (Graph attention and Graph transformer) architectures.

## CODE AND DATA AVAILABILITY

We did not generate any new dataset for this study but rather downloaded the reference datasets from the reference resources^15,36^. The source codes of EpiExpr-1D and EpiExpr-3D are hosted in GitHub (https://github.com/souryacs/3CExpr) along with the corresponding documentation. Snakemake pipelines to create the training datasets for both of these models are also provided within the same repository. The source code for the adapted Epi-GraphReg model, for application in different cell types, is hosted in GitHub (https://github.com/souryacs/EpiGraphReg_Custom).

---

**Supplementary Figure 1:**
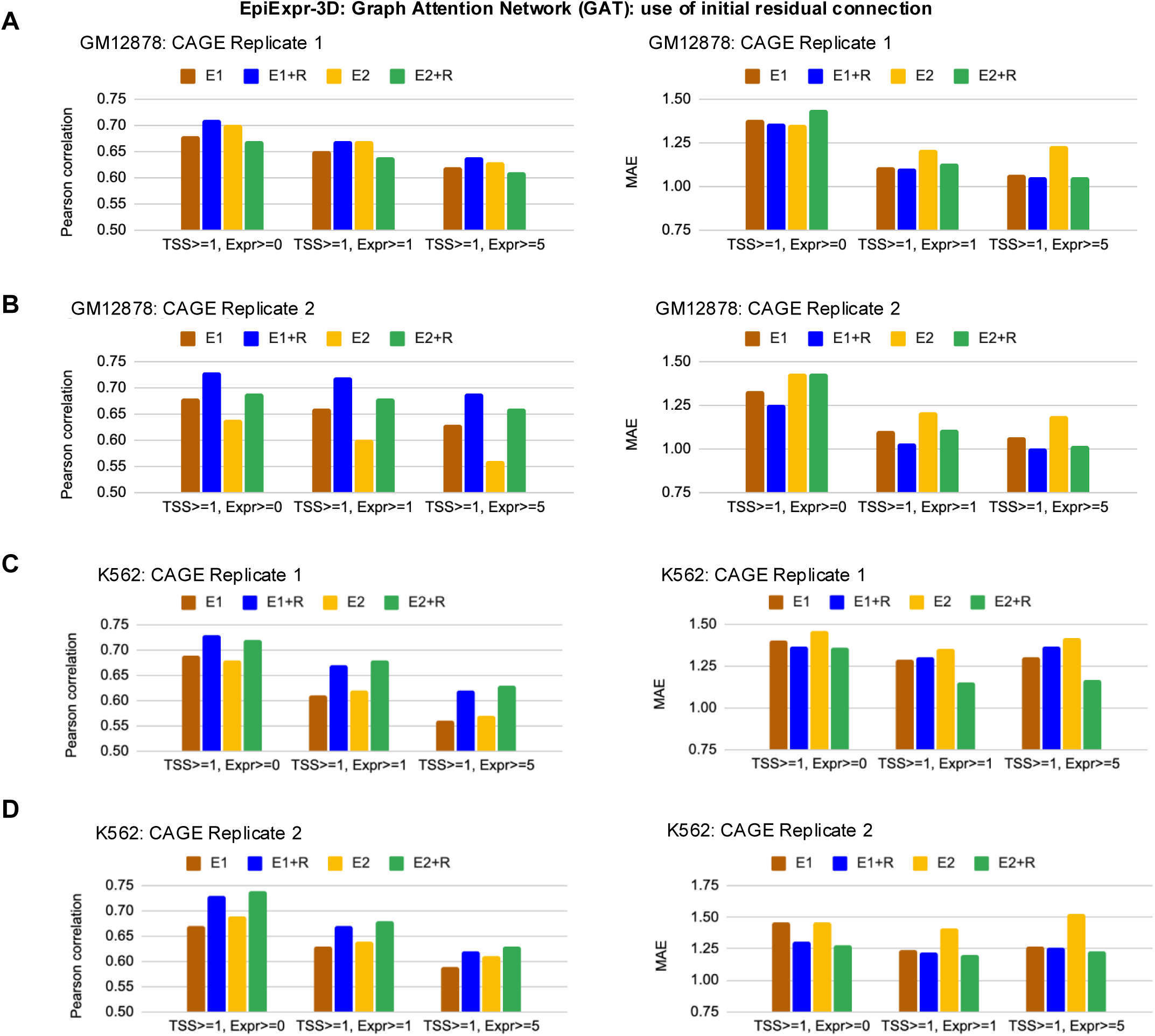
Comparison between different EpiExpr-3D settings with Graph Attention Network (GAT), different edge normalization techniques (E1 or E2) and using residual connection (R) or not, with respect to the GM12878 and K562 datasets from the Epi-GraphReg study. **(A)** Comparison of Pearson correlation (left) and mean absolute error (MAE) (right) between the true and predicted expressions for different settings of EpiExpr-3D with GAT, with respect to the epigenomic datasets and CAGE track replicate 1 for GM12878 cell type provided in the Epi-GraphReg paper. Track resolutions: epigenetic 100 bp, CAGE 5Kb. Validation chr: 1, 11; test chr: 2, 12. TSS>= *x,* Expr >= *y*: expression bins with at least *x* number of overlapping TSS and minimum expression *y*. E1: scikit-learn row normalization of graph edges, E2: double stochastic normalization of graph edges, R: using initial residual connection on GAT. **(B-D)** Similar comparisons for GM12878 and K562 cell types, with respect to different CAGE tracks.

**Supplementary Figure 2:**
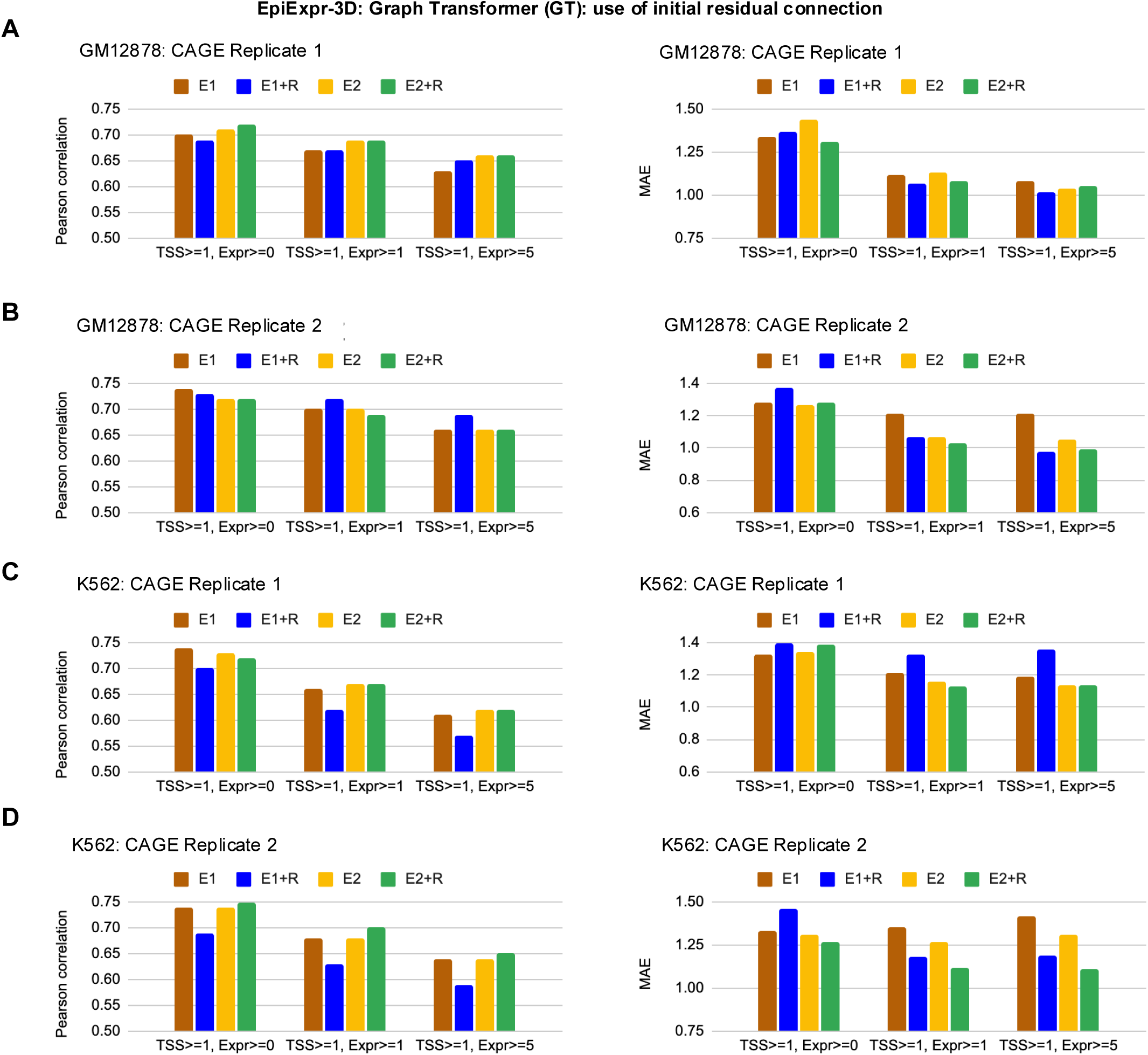
Comparison between different EpiExpr-3D settings with Graph Transformer (GT), different edge normalization techniques (E1 or E2) and using residual connection (R) or not, with respect to the GM12878 and K562 datasets from the Epi-GraphReg study. **(A)** Comparison of Pearson correlation (left) and mean absolute error (MAE) (right) between the true and predicted expressions for different settings of EpiExpr-3D with GT, with respect to the epigenomic datasets and CAGE track replicate 1 for GM12878 cell type from the Epi-GraphReg paper. Track resolutions: epigenetic 100 bp, CAGE 5Kb. Validation chr: 1, 11; test chr: 2, 12. TSS>= *x,* Expr >= *y*: expression bins with at least *x* number of overlapping TSS and minimum expression *y*. E1: scikit-learn row normalization of graph edges, E2: double stochastic normalization of graph edges, R: using initial residual connection on GT. **(B-D)** Similar comparisons for GM12878 and K562 cell types, with respect to different CAGE tracks.

**Supplementary Figure 3:**
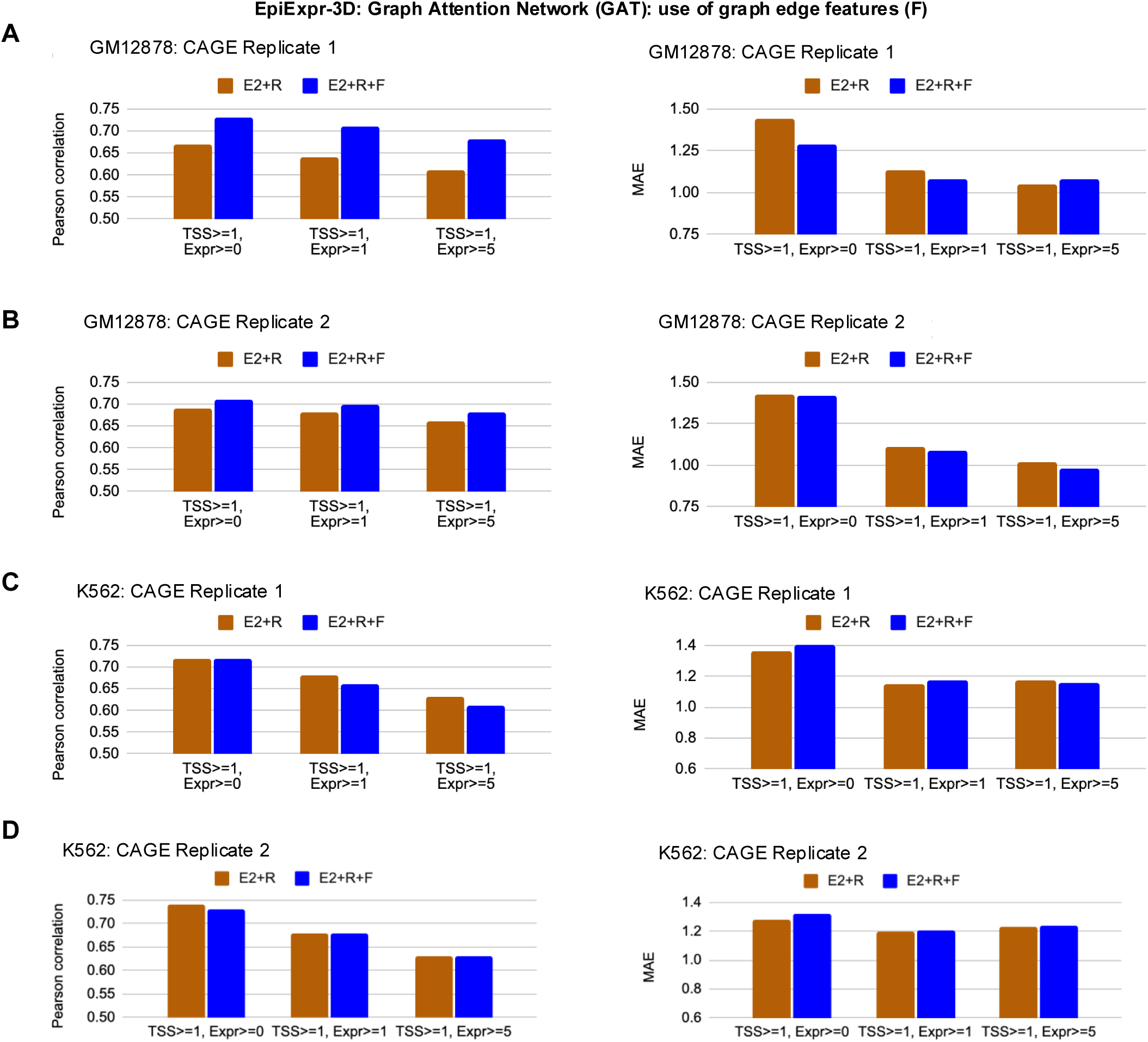
Comparison between EpiExpr-3D with GAT, without or with employing normalized chromatin contacts as graph edge features (indicated by the setting F), with respect to the GM12878 and K562 datasets from the Epi-GraphReg study. **(A)** Comparison of Pearson correlation (left) and mean absolute error (MAE) (right) between the true and predicted expressions for EpiExpr-3D with different settings of GAT, with respect to the epigenomic datasets and CAGE track replicate 1 for GM12878 cell type from the Epi-GraphReg paper. Various GAT settings with double stochastic edge normalization (E2), initial residual connection (R), and without (E2+R) or with normalized chromatin contact counts as graph edge features (E2+R+F) were tested. Track resolutions: epigenetic 100 bp, CAGE 5Kb. Validation chr: 1, 11; test chr: 2, 12. TSS>= *x,* Expr >= *y*: expression bins with at least *x* number of overlapping TSS and minimum expression *y*. E1: scikit-learn row normalization of graph edges, E2: double stochastic normalization of graph edges, R: using initial residual connection on GAT, F: using graph edge features. **(B-D)** Similar comparisons for GM12878 and K562 cell types, with respect to different CAGE tracks.

**Supplementary Figure 4:**
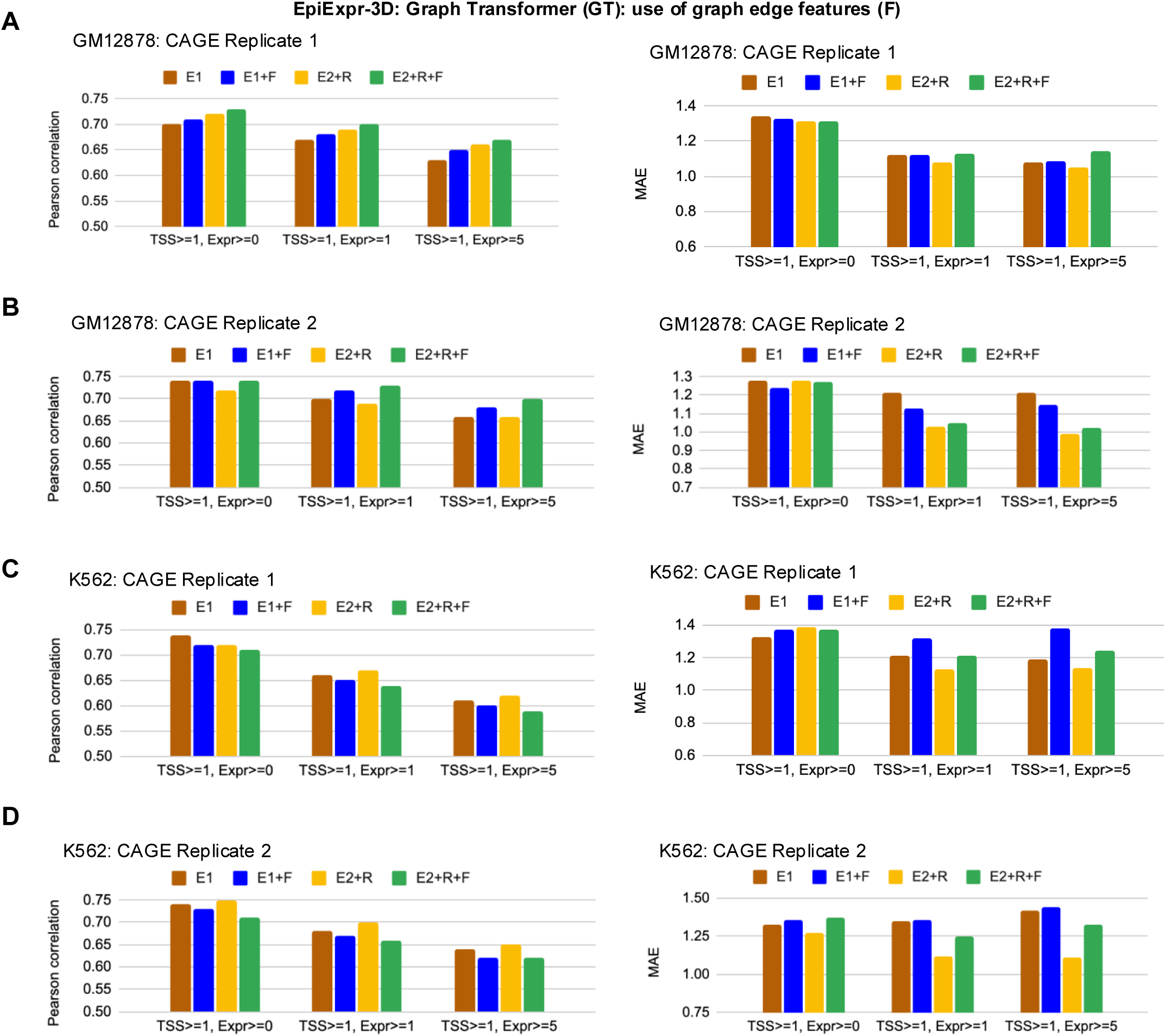
Comparison between EpiExpr-3D with Graph Transformer (GT), without or with employing normalized chromatin contacts as graph edge features (indicated by the setting F), with respect to the GM12878 and K562 datasets from the Epi-GraphReg study. **(A)** Comparison of Pearson correlation (left) and mean absolute error (MAE) (right) between the true and predicted expressions for EpiExpr-3D with different settings of GT, with respect to the epigenomic datasets and CAGE track replicate 1 for GM12878 cell type from the Epi-GraphReg paper. Various GT settings with either row-normalization of graph edges (E1) or double stochastic edge normalization (E2), without or with initial residual connection (R), and without or with normalized chromatin contact counts as graph edge features (F), have been benchmarked. Track resolutions: epigenetic 100 bp, CAGE 5Kb. Validation chr: 1, 11; test chr: 2, 12. TSS>= *x,* Expr >= *y*: expression bins with at least *x* number of overlapping TSS and minimum expression *y*. E1: scikit-learn row normalization of graph edges, E2: double stochastic normalization of graph edges, R: using initial residual connection on GT, F: using graph edge features. **(B-D)** Similar comparisons for GM12878 and K562 cell types, with respect to different CAGE tracks.

**Supplementary Figure 5:**
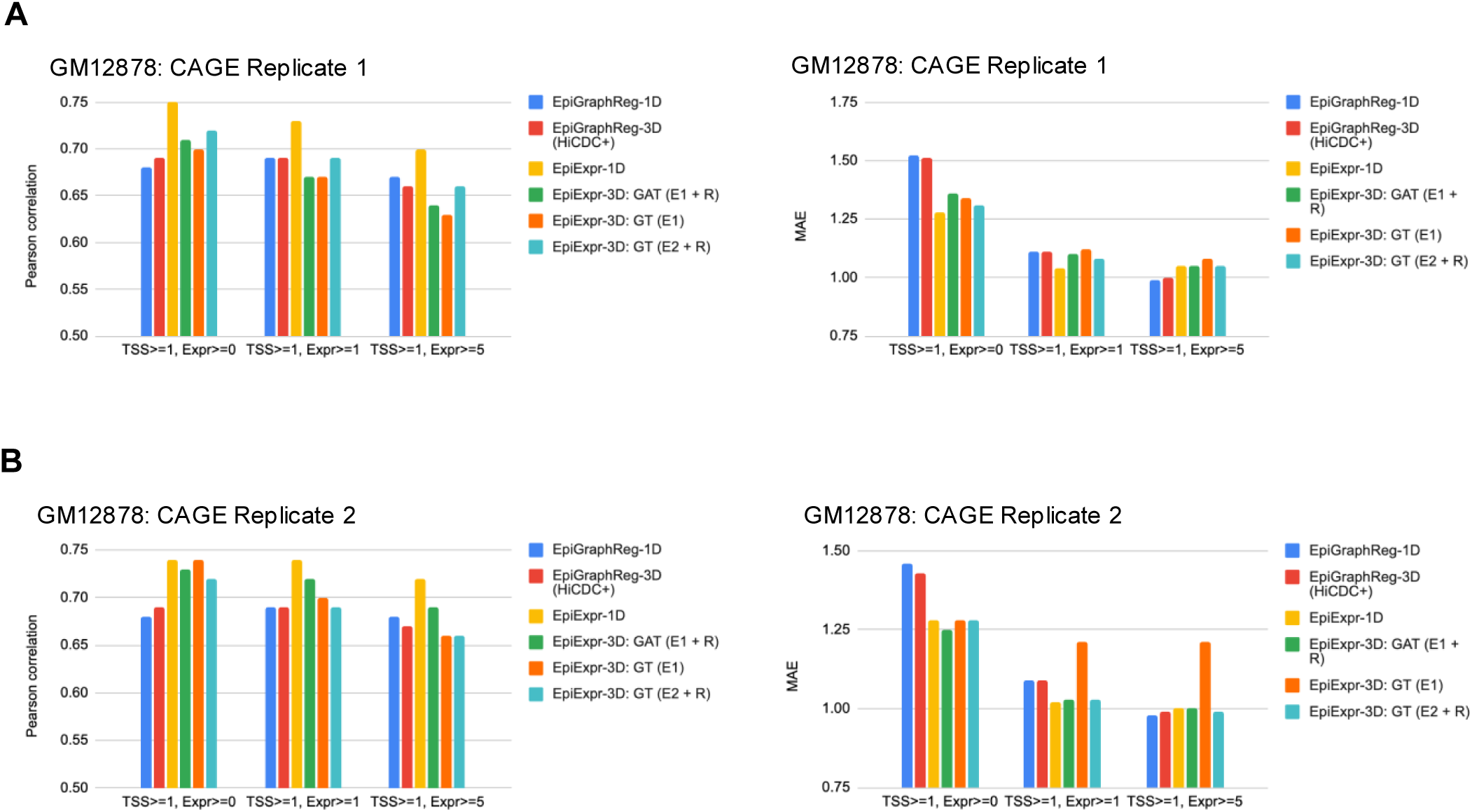
Performance comparison between EpiExpr-3D and Epi-GraphReg-3D with respect to the GM12878 datasets from the Epi-GraphReg study. **(A)** Comparison of Pearson correlation (left) and mean absolute error (MAE) (right) between the true and predicted expressions for Epi-GraphReg (1D and 3D with HiCDC+ loops) and different settings of EpiExpr-3D, with respect to the epigenomic datasets and CAGE track replicate 1 for GM12878 cell type from the Epi-GraphReg paper. Track resolutions: epigenetic 100 bp, CAGE 5Kb. Validation chr: 1, 11; test chr: 2, 12. TSS>= *x,* Expr >= *y*: expression bins with at least *x* number of overlapping TSS and minimum expression *y*. GAT: graph attention network, GT: graph transformer, E1: scikit-learn row normalization of graph edges, E2: double stochastic normalization of graph edges, R: using initial residual connection on GAT / GT. **(B)** Similar to (A) for another CAGE track of the GM12878 cell type.

